# New PALM-compatible integration vectors for use in the Gram-positive model bacterium *Bacillus subtilis*

**DOI:** 10.1101/2023.03.16.532899

**Authors:** Ipek Altinoglu, Rut Carballido-Lopez

**Affiliations:** MICALIS Institute, INRAE, AgroParisTech, Université Paris-Saclay, 78350 Jouy-en-Josas, France

**Keywords:** *Bacillus subtilis*, PALM, integration vector, dronPA, PAmCherry, LacI, bacteriophage SPP1

## Abstract

Improvements in super-resolution and single-molecule techniques together with the development of new fluorescent proteins and labelling methods have allowed super-resolution microscopy to be applied to bacterial cells. Cloning vectors remain important tools for researchers to perform efficient labelling. Here, we describe the creation of four PhotoActivated Localization Microscopy (PALM)-compatible integration plasmids for the Gram-positive model organism *Bacillus subtilis*. These plasmids carry either the photoswitchable green fluorescent protein dronPA or the photoactivatable red fluorescent protein PAmCherry1, codon-optimized or not for *B. subtilis*. For fast and interchangeable cloning, we have inserted multi cloning sites at both the C-terminal and the N-terminal end of the fluorophores. The plasmids replicate in *Escherichia coli* and allow integration at the ectopic *amyE* or *thrC* loci of *B. subtilis* via double homologous recombination, for stable chromosomal insertions of dronPA and PAmCherry1 protein fusions respectively. Dual color imaging is accessible with the simultaneous use of the two vectors. Insertion of the gene encoding the LacI repressor under control of a constitutive promoter in each of the four plasmids yielded four derivative vectors that, combined with an array of *lacO* operator sites, allow Fluorescent Repressor-Operator System (FROS) localization studies. We demonstrated the successful photoactivation of the LacI-dronPA or PAmCherry1 fusions, and used them to report with nanoscale precision the subcellular localization of bacteriophage SPP1 DNA in infected *B. subtilis* cells as proof of concept. Our PALM-compatible integration vectors expand the genetic toolbox for single molecule localization microscopy studies in *B. subtilis*.

## INTRODUCTION

Over the last two decades, fluorescence light microscopy has become a key tool for understanding the cellular and subcellular organization of bacterial cells [1,2]. Despite the low-cost and rapid solutions obtained with conventional light microscopies, their resolution remains limited by the diffraction of light (Abbe’s diffraction limit), typically to ∼200-300 nm laterally and ∼500-700 nm axially in biological specimens [3]. Super-resolution (SR) microscopy techniques allow to overcome this limitation [4-8] and have become available for prokaryotic cell biology studies over the last decade. SR techniques allow to resolve subcellular structures and provide a better understanding of relative positions of tagged proteins, DNA, RNA as well as their interactions and dynamic localizations with nanoscopic precision. They can be classified in two groups depending on their working principle, which can be deterministic or stochastic. The main difference between the two groups is the protein labelling method. In deterministic SR techniques such as structured illumination microscopy (SIM) and stimulated emission depletion (STED) microscopy, the proteins of interest are labelled with conventional fluorophores (fluorescent proteins (FP) or chemical dyes) that show a non-linear response to the excitation used to enhance resolution [5,6]. In contrast, stochastic SR techniques require specific fluorophores that upon activation with the adequate laser allow stochastic switching on and off of single-molecule fluorescence signals over time, allowing the so-called single molecule localization microscopy (SMLM). SMLM-based techniques include stochastic optical reconstruction microscopy (STORM), point accumulation in nanoscale topography (PAINT) and photo activated localization microscopy (PALM) [4,7,8]. In PALM, specific fluorophores are used to activate and image single molecules allowing quantitative analysis and fine-scale localizations with lateral resolution down to ∼10–30 nm [9]. Three types of FPs are commonly used for PALM imaging: (a) photoactivatable, which emit light upon activation with UV light (e.g. PA-GFP, PAmCherry1, [10,11]); (b) photoconvertible, which irreversibly change their emission spectrum (most form green to red) upon activation with UV light (e.g. mEos3.2, mMaple3 [12,13]) or (c) photoswitchable, which repeatedly switch between non fluorescent (‘dark’) and fluorescent states upon UV activation, until photobleaching (e.g. dronPA [14]). Using the right fluorophores and assessing the expression levels, the functionality and the stability of FP fusions is crucial in PALM imaging. The FP tag or overexpression of the FP fusion can affect the function and/or the organization of the protein of interest. Oligomerization of FPs expressed in high concentration can also lead to artefactual localization patterns and clusters, which prompted for the development of monomeric FPs [15]. Furthermore, export of bacterial envelope proteins via the general bacterial secretion system (Sec) in an unfolded conformation, prompted for the use of FPs with enhanced folding and chromophore formation (maturing) kinetics and with increased stability, such as superfolding derivatives of GFP (sfGFP, [16]) for periplasmic and extracellular protein localization studies [17]. Finally, codon-optimisation of the FP can improve their expression and brightness in heterologous bacterial systems.

*Bacillus subtilis* is a rod-shaped, aerobic soil bacterium [18]. It is so far the best-characterized Gram-positive bacterium for several reasons such as high genetic tractability, survival in extreme conditions by cellular differentiation to form highly resistance endospores, ease to grow and manipulate in laboratory conditions and efficiency in enzyme production for industrial use [19- 22]. It has been used for decades as model organism for cell biology and subcellular organization studies in bacteria, to study chromosome segregation and DNA replication, cell cycle, cell morphogenesis, biofilm formation, sporulation, viral infection and antibiotic resistance, among other cellular processes [23-26]. The ease of genetic manipulation of *B. subtilis* results from the combination of its natural competence for genetic transformation and its highly efficient homologous recombination. These features allow integration vectors instead of replicative vectors to be commonly used due to their stable maintenance (even in the absence of selective pressure) when inserted in the chromosome via double homologous recombination, and to the defined single copy number of the integrated genes [27-31]. Vectors (here restricted to plasmid vectors) have been therefore extensively modified to meet custom needs and applications. Four nonessential integration loci, *thrC, amyE, sacA* and *lacA*, are traditionally used for ectopic insertion of genetic constructions in *B. subtilis*. Here, we have developed and tested four integrative vectors, two for insertion into the *thrC* locus and two into the *amyE* locus, to express fusions to the photoactivatable monomeric red FP PAmCherry1 or to PAmCherry1 codon-optimized for expression in *B. subtilis* (bsPAmCherry1) and to the photoswitchable green FP dronPA or to dronPA codon-optimized for expression in *B. subtilis* (bsdronPA), respectively. We confirmed the integration of the vectors and the functionality of the fluorophores in both live and fixed cells using the commonly used Fluorescent Repressor-Operator System (FROS) LacI*-lacO* [32,33]. Bacteriophage SPP1 DNA, engineered to carry an array of ∼64 *lacO* operator sites inserted in its genome, was previously visualized during infection of *B. subtilis* cells producing a chromosomally-encoded LacI-mCherry fusion [34,35]. We constructed four derivative vectors that allow *lacI-PAmCherry1* or *lacI- dronPA*, or their codon-optimized versions, expressed under control of a constitutive promoter, to be integrated into the *B. subtilis* chromosome and used them to reveal the nanoscale localization of bacteriophage SPP1 DNA by single color PALM as proof of concept.

## MATERIALS AND METHODS

### Bacterial strains and growth conditions

The bacterial strains and phage used in this study are listed in Table 1. *B. subtilis* YB886 [36] and *E. coli* DH5α were used as wild-type strains. Unless specified otherwise, bacterial strains were grown in lysogeny broth (LB) or LB agar (1.5 %) and, when appropriate, antibiotics were used at the following concentrations: ampicillin 100 μg/mL, chloramphenicol 25 μg/mL, erythromycin 30 μg/mL for *E. coli* and chloramphenicol 5 μg/mL, erythromycin 0.5 μg/mL, lincomycin 12.5 μg/mL, streptomycin 100 μg/mL for *B. subtilis*.

**Table 1.** Bacterial strains and phage used in this study.

| Strain name | Strain type | Description | Reference |
| --- | --- | --- | --- |
| Bacterial strains |  |  |  |
| DH5α | *E.coli* | competent cells | Lab stock |
| ECRCL51 | *E.coli* | DH5α/pDG364 | Lab stock |
| ECRCL67 | *E.coli* | DH5α/pDG1664 | Lab stock |
| ECRCL0226 | *E.coli* | DH5α/pIA019 | Eurofins |
| ECRCL0227 | *E.coli* | DH5α/pIA020 | Eurofins |
| ECRCL0287 | *E.coli* | DH5α/pIA021 | This study |
| ECRCL0289 | *E.coli* | DH5α/pIA023 | This study |
| ECRCL0290 | *E.coli* | DH5α/pIA025 | This study |
| ECRCL0292 | *E.coli* | DH5α/pIA027 | This study |
| ECRCL0293 | *E.coli* | DH5α/pIA028 | This study |
| ECRCL0295 | *E.coli* | DH5α/pIA030 | This study |
| ECRCL0297 | *E.coli* | DH5α/pIA025 | This study |
| ECRCL0299 | *E.coli* | DH5α/pIA034 | This study |
| ECRCL0300 | *E.coli* | DH5α/pIA033 | This study |
| ECRCL0301 | *E.coli* | DH5α/pIA035 | This study |
| ECRCL0302 | *E.coli* | DH5α/pIA036 | This study |
| YB886 | *B. subtilis* | amyE trpC2 metB5 xin-1 attSPβ | [36] |
| GSY10004 | *B. subtilis* | YB886, thrC::(*P*<sub>pen</sub>-lacIΔ11-mcherry mls erm2) | [39] |
| RCL0937 | *B. subtilis* | YB886, amyE::(*P*<sub>pen</sub>-lacIΔ11-dronPA mls cm) | This study |
| RCL0938 | *B. subtilis* | YB886, thrC::(*P*<sub>pen</sub>-lacIΔ11-PAmCherry1 mls erm2) | This study |
| RCL0939 | *B. subtilis* | YB886, thrC::(*P*<sub>pen</sub>-lacIΔ11-bsPAmCherry1 mls erm2) | This study |
| RCL0941 | *B. subtilis* | YB886, amyE::(*P*<sub>pen</sub>-lacIΔ11-bsdronPA mls cm) | This study |
| Phage |  |  |  |
| lacO64 | SPP1 | SPP1delX110lacO64 | [35] |

### DNA manipulation and plasmids construction

#### General cloning procedures

Polymerase chain reaction (PCR) amplifications were carried out using the Phusion^®^ polymerase. Restriction enzyme digestions were performed using New England Biolabs (NEB) high fidelity enzymes with the protocol suggested by the supplier. All plasmids were constructed by isothermal assembly [37], using a minimum of 20 bp overlapping regions between DNA fragments with custom made kit Plasmid selection and verification after the transformation were checked using the OneTaq^®^ 2X Master Mix with Standard Buffer (NEB). Plasmids were purified using the Qiagen plasmid purification kit according to manufacturer’s protocol.

The plasmids and oligonucleotides used in this study are listed in Table 2 and Table 3, respectively.

**Table 2.** List of plasmids.

| Plasmid number | Description | Reference |
| --- | --- | --- |
| pDG364 | *amyE::cm* | [45] |
| pDG1664 | *thrC::erm2 (thrC Amp<sup>r</sup> Erm<sup>r</sup> Spec<sup>r</sup>)* | [27] |
| pPT300 | *thrC::(P<sub>pen</sub>-lacIΔ11-mcherry mls erm)* | [39] |
| pEYY133 | Cloning vector for C terminal *PAmCherry1* fusion | [38] |
| pEYY99 | Cloning vector for C terminal *dronPA* fusion | [38] |
| pIA019 | *pEX-A128-bsdronPA* | This study |
| pIA020 | *pEX-A128-bsPAmCherry1* | This study |
| pIA021 | *amyE::(MCS cm)* | This study |
| pIA023 | *thrC::(MCS erm2)* | This study |
| pIA025 | *thrC::(P<sub>pen</sub>-lacIΔ11-bsPAmCherry1 mls erm2)* | This study |
| pIA027 | *amyE::(dronPA cm), for dronPA fusions* | This study |
| pIA028 | *thrC::(PAmCherry1 erm2), for PAmCherry1 fusions* | This study |
| pIA030 | *thrC::(P<sub>pen</sub>-lacIΔ11-PAmCherry1 erm2)* | This study |
| pIA033 | *thrC::(bsPamCherry1 erm2), for bsPAmCherry1 fusions* | This study |
| pIA034 | *amyE::(bsdronPA cm), for bsdronPA fusions* | This study |
| pIA035 | *amyE::(P<sub>pen</sub>-lacIΔ11-dronPA cm)* | This study |
| pIA036 | *amyE::(P<sub>pen</sub>-lacIΔ11-bsdronPA cm)* | This study |

**Table 3.**
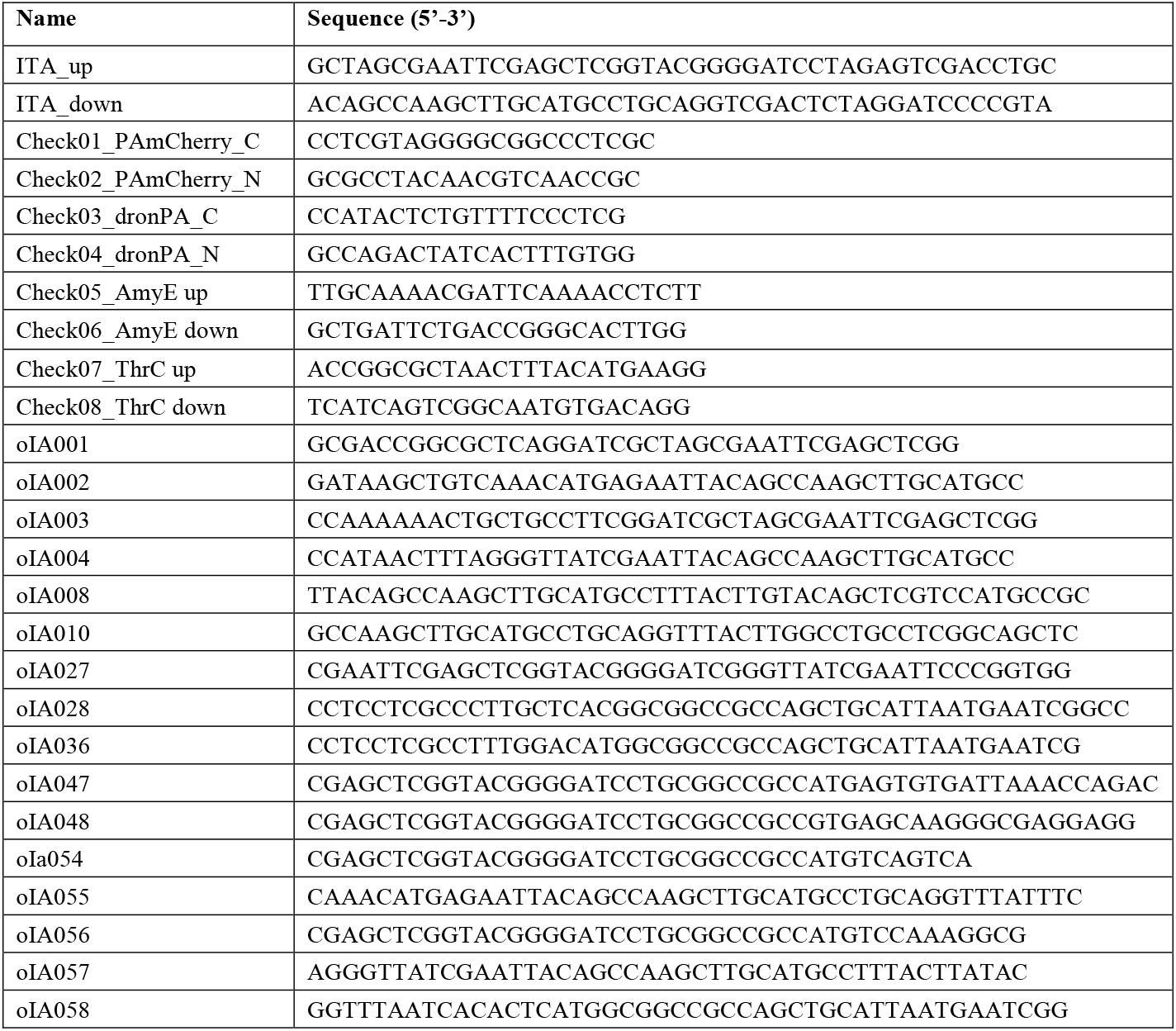
List of oligonucleotides.

#### Construction of Multi Cloning Sites (MCS)

The overlapping oligonucleotides ITAup (45 bp) and ITAdown (45 bp) were annealed by DNA polymerase reaction. The resulting double-stranded DNA is called MCS (Figure 1B).

**Figure 1.**
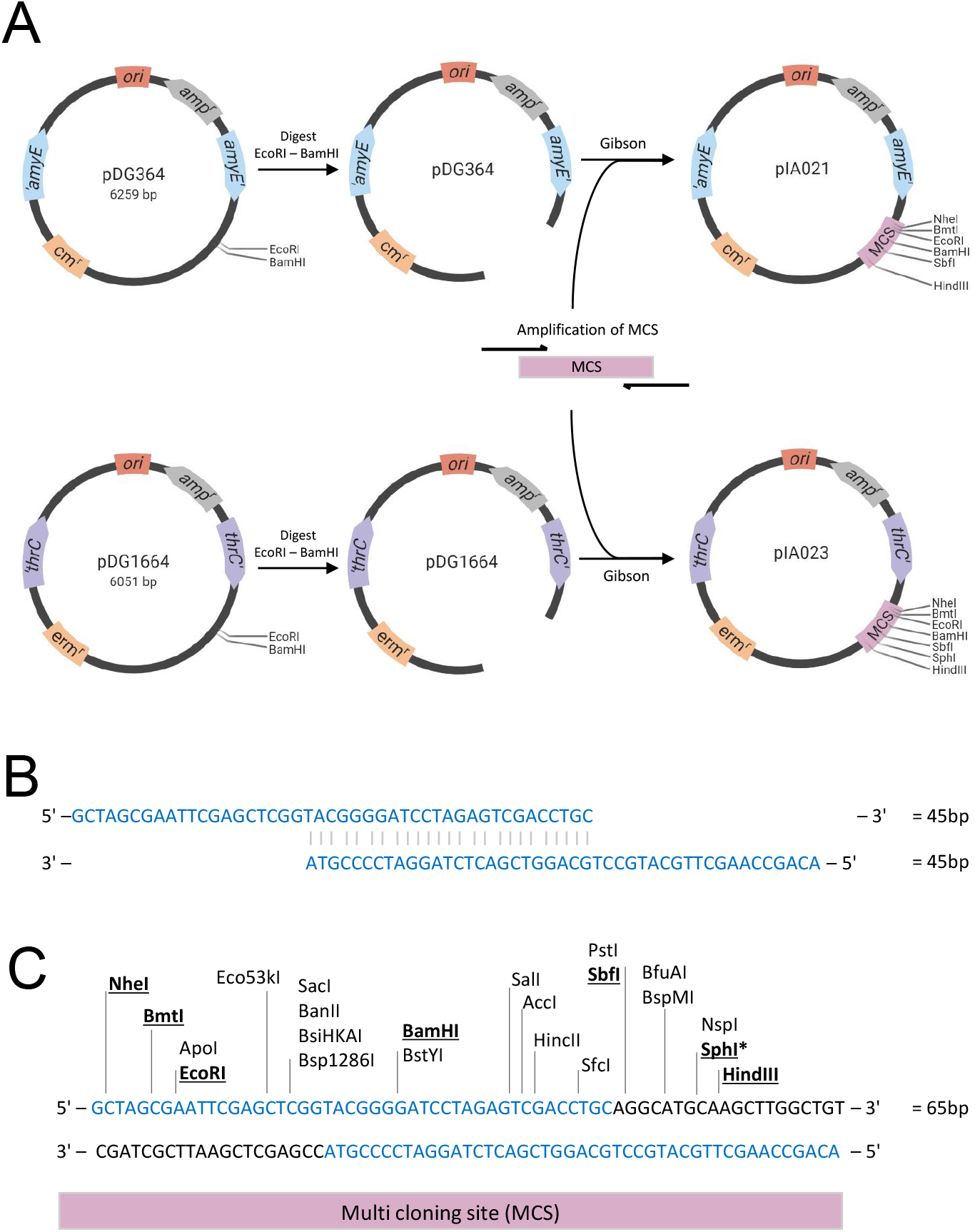
Building empty integrative plasmids for efficient cloning. **A**. Plasmids pDG364 and pDG1664 were used as backbones. The replication origin (*ori*) and ampicillin resistance cassette (amp^r^) for vector verification and propagation in *E. coli* are marked in light red and gray, respectively. The *B. subtilis* genomic integration sites are shown in blue for *amyE* and in purple for *thrC*. The chloramphenicol and erythromycin resistance cassettes (cm^r^ and erm^r^) for selection in *B. subtilis* are indicated in orange. The multi cloning site (MCS) is shown in pink. Arrows indicate the plasmid construction process (see the material and methods section for details). The MCS was inserted into the pDG364 and pDG1664 empty plasmids by Gibson Assembly. **B-C**. The MCS was engineered using two overlapping oligonucleotides of 45 bp each (B) that were annealed to generate a double stranded DNA fragment of 65 bp (C) containing 22 restriction sites. Underlined in bold are the restriction sites unique in the new plasmids pIA021 and pIA023. The star indicates the SphI site, which is only unique in the MCS for plasmid pIA023.

#### Construction of empty integration vectors pIA021 and pIA023

MCS was amplified using primers oIA001 and oIA002 and cloned through Gibson Assembly into pDG364 [30] (*amyE::cm*) digested with EcoRI and BamHI, resulting in plasmid pIA021. MCS amplified with primers oIA003 and oIA004 was cloned into pDG1664 [27] (*thrC::erm2*) digested by EcoRI and BamHI, resulting in primer pIA023.

#### Construction of plasmids pIA027, pIA034, pIA028 and pIA033, containing PALM-compatible fluorescent proteins

PALM-compatible fluorophores dronPA (in plasmid pEYY99) and PAmCherry1 (in plasmid pEYY133) were kindly provided by Dr. Yoshiharu Yamaichi [38]. The *dronPA* and *PAmCherry1* genes were sequenced for confirmation. The *dronPA* and *PAmCherry1* gene sequences codon- optimized for *B. subtilis* (*bsdronPA* and *bsPAmCherry1)* were synthesized and cloned into a pEX backbone plasmid by Eurofins Genomics, giving pIA019 (pEX-A128-bsdronPA) and pIA020 (pEX-A128-bsPAmCherry1).

The *dronPA* sequence was amplified from pEYY99 using primers oIA010 and oIA047, then cloned into pIA021 digested with BamHI and SbfI, resulting in plasmid pIA027, *amyE::(dronPA cm)*. The *PAmCherry1* sequence was amplified from pEYY133 using primers oIA008 and oIA048, then cloned into pIA023 digested with BamHI and SphI, resulting in plasmid pIA028, *thrC::(PAmCherry1 erm2). bsdronPA* was amplified from pIA019 using primers oIA054 and oIA055, then cloned into pIA021 digested with BamHI and HindIII, giving plasmid pIA034, *amyE::(bsdronPA cm). bsPAmCherry1* was amplified from pIA020 using primers oIA056 and oIA057, then cloned into pIA023 digested with BamHI and HindIII, giving plasmid pIA033, *thrC::(bsPamCherry1 erm2)*.

All plasmids were verified by Sanger DNA sequencing (GATC Biotech/Eurofins Genomics, Ebersberg, Germany). Plasmids *pIA027, pIA034, pIA028 and pIA033* were fully sequenced and deposited at Addgene (plasmid catalog numbers pending).

#### Construction of plasmids pIA035, pIA036, pIA030 and pIA025, containing PALM-compatible fluorescent fusions to LacI

Plasmid pPT300, *thrC::(Ppen-lacI*Δ*11-mcherry mls erm)* was kindly provided by Dr. Paulo Tavares [39]. The *lacI* ORF together with the RBS and promoter region sequences were amplified and inserted in plasmids pIA027, pIA034, pIA028 and pIA033 as follows.

pPT300 was amplified using primers oIA027 and oIA058. The resulting fragment was cloned into pIA027 digested by BamHI and NotI, giving plasmid pIA035, *amyE::(Ppen-lacI*Δ*11-dronPA cm)*. pPT300 was amplified using primers oIA027 and oIA069. The resulting fragment was cloned into pIA034 digested with BamHI and NotI, giving pIA036, *amyE::(Ppen-lacI*Δ*11-bsdronPA cm)*. pPT300 was amplified using primers oIA027 and oIA028. The resulting fragment was cloned into pIA028 digested with BamHI and NotI, giving pIA030, *thrC::(Ppen-lacI*Δ*11-PAmCherry1 erm2)*. pPT300 was amplified using primers oIA027 and oIA036. The resulting fragment was cloned into pIA033 digested with BamHI and NotI, giving pIA025, *thrC::(Ppen-lacI*Δ*11-bsPAmCherry1 mls erm2)*.

All constructs were verified by Sanger DNA sequencing (GATC Biotech/Eurofins Genomics, Ebersberg, Germany). Plasmids *pIA035, pIA036, pIA030 and pIA025* plasmids were then fully sequenced and deposited at Addgene (plasmid catalog numbers pending).

### Transformation and selection

*E. coli* DH5α competent cells were used for the routine cloning applications and transformation of all plasmids at minimum 50 ng/μl. Cells harboring the plasmids were selected by plating transformants on LB agar plates containing the appropriate antibiotic and incubated overnight at 37ºC. Single colonies grown overnight on the selective medium were screened by colony PCR to determine the presence or absence of the fragment inserted in the plasmid using check-primers between Check01 and Check08 in Table 3. Positive colonies were grown overnight in LB medium at 37ºC with agitation (180 rpm) and plasmids were purified.

*B. subtilis* YB886 competent cells were used for integration of the FROS vectors pIA035, pIA036, pIA030 and pIA025. Competent cells were prepared by a two-step starvation protocol as previously described [40]. Minimum 1 μg of circular plasmid was added to 500 μl of competent cells. Cells were plated onto appropriate selective media and incubated overnight at 37°C. Single colonies grown overnight were patched onto replica plates for replication prior to use in colony PCR to amplify *amyE* or *thrC* regions.

### Sample preparation for fluorescence microscopy

*B. subtilis* strains were plated with appropriate antibiotic selection and incubated overnight at 37ºC to isolate single colonies. A single colony was grown overnight in LB medium at 30ºC with agitation (180 rpm). Overnight cultures were diluted 1:100 in fresh LB medium and grown at 37ºC with agitation (180 rpm). When cultures reached an OD_600nm_ of 0.8, they were supplemented with 10 mM CaCI_2_ and infected with phage SPP1 (strain SPP1*delX110lacO*_*64*_) at an input multiplicity of 1. Infected cells were further incubated at 37ºC with orbital agitation (180 rpm) for 50 min and spotted onto an agarose pad (1% agarose with H_2_O) mounted on a glass slide using a Gene Frame (Thermofisher). Cover slips (thickness No. 1.5H) and/or slides were pre-cleaned with acetone and then Plasma Cleaned with Harrick Plasma (PDC-002-CE 230V) for 10 minutes, high level.

Image acquisitions of live cells were performed at 50 min post-infection (p.i.). For image acquisition of fixed cells, infected cells were fixed 50 min p.i. with fresh fixing solution (1 M KPO_4_ pH7, 2% PFA, 0.02% Glutaraldehyde) in a 2:1 ratio (2 vol. cells:1 vol. solution). Fixed cells incubated were 15 min at RT, then 30 min in ice, washed with 100 μl of PBS and kept overnight at 4ºC.

All images were acquired on a Zeiss Elyra PS1 microscope, at 37ºC for live cells and at RT for fixed cells. The Zeiss Elyra PS1 was equipped with a 63x (NA 1.40) and an Apo 100x (NA 1.46) Apochromatic oil immersion objectives, coherent lasers emitting at 405 nm (50 mW), 488 nm (100 mW) and 561 nm (100 mW), an emCCD Andor iXon 897 camera and a PCO edge sCMOS camera. The microscope and cameras were controlled by the Zen software version ZEN 2012 SP2.

### Photoactivated Localization Microscopy (PALM) imaging

PALM images were taken in fields of 512 × 512 pixels using the emCCD camera, 100x (NA 1.46) objective, a HILO angle 43.74º and 25 ms exposures. The 488 or 561 nm lasers were used for the first 1 500 frames to bleach out cell background. Continued activation with the 405 nm and the 488/561 lasers was used for molecule detection over 1500^th^ -4000^th^ frames. The ThunderSTORM [41] plug-in of Fiji was used for molecule detection and quantification analysis.

### Structured Illumination Microscopy (SIM) imaging

SIM images were taken in fields of 256 × 256 pixels using the sCMOS camera and the 63x (NA 1.40) objective. 100 ms exposures were used for white field illumination and 50 ms exposures were used for the 561 nm laser at 50 %. 34.0 μm G3 grid with 3 phases and 5 rotations was used for SIM imaging. Re-construction of SIM images was made using the FairSIM [42] plug-in of Fiji [43] and the Zen 2012 SP2 image analysis software equipped for 2D.

## RESULTS

### Design of the integration vectors and insertion of Multi-Cloning-Sites

To generate standardized cloning plasmids for endogenous expression of PALM-compatible FP fusions upon double cross-over homologous recombination in *B. subtilis*, we chose as plasmid backbones two well-established empty vectors, pDG364 and pDG1664, which integrate at the *amyE* and *thrC* loci of the *B. subtilis* chromosome, respectively [27,30] (Figure 1A). Both plasmids contain the *E. coli* origin of replication (*ori*), to maintain and propagate the foreign plasmid in *E. coli*, and the *bla* gene (indicated as AmpR in the plasmids maps), which encodes β-lactamase and thus allows the selection of the plasmid with ampicillin in *E. coli*. In addition, pDG364 and pDG1664 harbor *cm* and *erm* cassettes, respectively, for selection in *B. subtilis*. We chose two different integration loci and selection markers to allow the final vectors to be combined in the same strain for multicolor imaging. To facilitate the cloning, we engineered a Multi-Cloning-Site (MCS) by using two overlapping oligonucleotides of 45 bp each (Figure 1B) that were annealed to generate a double stranded DNA fragment of 65 bp called MCS that includes 22 unique restriction sites (Figure 1C). Isothermal assembly (Gibson) [37] was used to integrate the MCS into pDG364 and pDG1662 linearized by EcoRI and BamHI digestion (Figure 1A). The resulting circular vectors pIA021 and pIA023 possess six common unique restriction sites: NheI, BmtI, EcoRI, BamHI, SbfI, HindIII, and pIA023 possesses one additional unique restriction site: SphI.

### New plasmids with PALM-compatible fluorescent proteins PAmCherry1 and dronPA

PALM-compatible FP play a critical role in SMLM image acquisition depending on their activation principle, photon detection, lifetime of the fluorophore and oligomerization properties. Beside single-color imaging, choosing the right combination of FP for multi-color imaging becomes even more complicated for PALM due to (*i*) the availability of laser sources, as two lasers at least (for activation and imaging) are needed for each fluorophore, and (*ii*) photoproperties of FPs, preventing photoconvertible fluorophores (e.g. mEos3.2, mMaple3), which use 3 laser sources, to be combined with any other PA FPs. Thus, it may take some serious time to define the most suitable fluorophore(s) for an experiment. To generate plasmids compatible with two-color imaging, we first chose a suitable FP pair: the photoswitchable monomeric green fluorescent protein dronPA (originally isolated from *Echinophyllia sp. SC22;* fluorophore excitation 503 nm, emission 518 nm (green), photoactivation with UV-Violet, 405 nm) and the photoactivatable monomeric red fluorescent protein PAmCherry1 (derived from mCherry from *Discosoma sp*. fluorophore excitation 564 nm, emission 595 nm (red), photoactivation with UV-Violet, 405 nm). The dronPA and PAmCherry1 are both initially in dark state, activated with 405 nm and spectrally distant ensuring minimal crosstalk. Furthermore, they are monomeric and have been successfully used for dual-color membrane labelling in bacterial cells [38].

We used plasmids pIA021 and pIA023 to introduce the photoswitchable FP dronPA and the photoactivatable FP PAmCherry1, respectively, into the MCS using isothermal assembly. Both the original version of the FP and a version codon-optimized for expression in *B. subtilis* were cloned. The resulting four new plasmids are pIA027 (*amyE::amyE-mcs-dronPA-mcs*), pIA034 (*amyE::amyE-mcs-bsdronPA-mcs*), pIA028 (*thrC::thrC-mcs-PAmCherry1-mcs*) and pIA033 (*thrC::thrC-mcs-bsPAmCherry1-mcs*) (Figure 2A and Table 2). They allow the cloning of the promotor region and of the protein of interest fused to the N-terminal of the FP by either isothermal assembly or classical enzyme digestion/ligation methods. Cloning of the protein of interest fused to the C-terminal of the FP is also possible by isothermal assembly. Pairwise combination of these plasmids can allow simultaneous insertion at *amyE* and *thrC* of two proteins of interest, fused to dronPA (or bsdronPA) and PAmCherry1 (or bsPAmCherry1) respectively, for dual-color PALM imaging.

**Figure 2.**
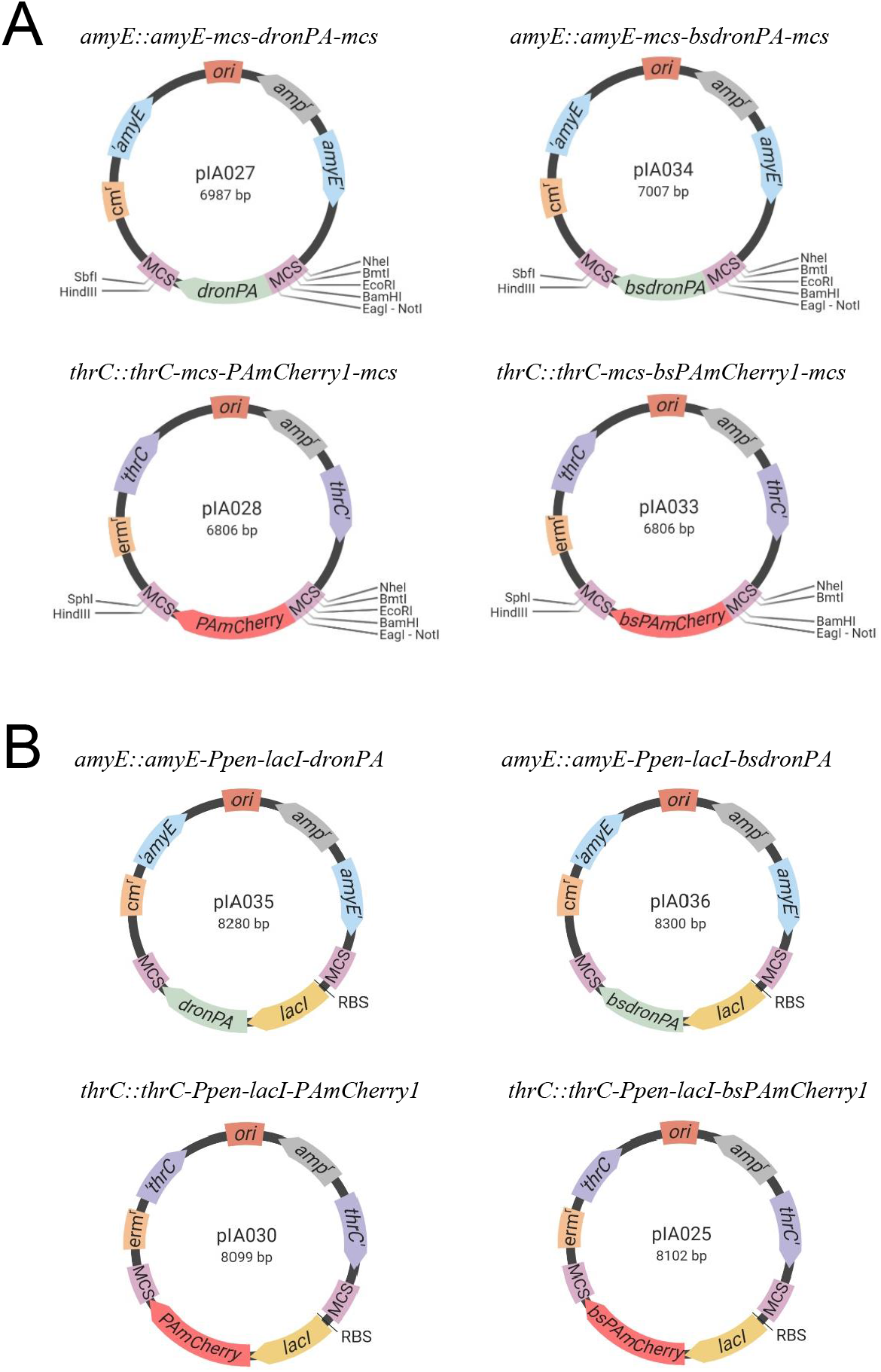
PALM-compatible integration plasmids for *B. subtilis*. The replication origin (*ori*) and ampicillin resistance cassette (amp^r^) for *E. coli* are marked in light red and gray, respectively. The *B. subtilis* genomic integration sites are shown in blue for *amyE* and purple for *thrC*, and the resistance cassettes are indicated in orange. The multi cloning sites (MCS) are shown is pink. The PALM-compatible fluorescent fusions are indicated in light green and light magenta for *dronPA*/*bsdronPA* and *PAmCherry1*/*bsPAmCherry*, respectively. **A**. Integration plasmids with the PALM-compatible fluorescent proteins dronPA and PAmCherry1. **B**. Integration plasmids for *lacO*/LacI FROS system in *B. subtilis*. The *lacI* gene, indicated in yellow, was cloned upstream the PALM-compatible fluorophores.

### Functionality of the plasmid-bore PALM-compatible fluorophores for FROS localization studies in *B. subtilis*

To confirm the functionality of our new vectors pIA027, pIA034, pIA028 and pIA033, we used a Fluorescent Repressor-Operator System (FROS) [32,33]. FROS are used to label specific DNA segments through binding of a fluorescently labelled repressor to repeated sequences of its cognate operator. They allow visualization of DNA loci in cells to understand genome organization and dynamics and to determine copy numbers of genetic loci [44]. Here, we chose to label bacteriophage SPP1 DNA using a *lacO*/LacI FROS system previously used to investigate the subcellular localization of the viral DNA during infection of *B. subtilis* [34,35]. Upon irreversible binding to the surface of a *B. subtilis* host cell, bacteriophage SPP1 injects its DNA into the cytoplasm of the bacterium, where the viral DNA replicates to produce viral proteins and assemble new virions, which are ultimately released in the medium upon lysis of the host cell [39]. Using a chromosomally-encoded, constitutively-expressed LacI-mCherry fusion and a phage engineered to carry an array of ∼64 *lacO* operator sites inserted in its genome (SPP1*delX110lacO64*), the viral DNA was visualized in infected cells, showing that a DNA compartment that harbors viral genome replication is formed and increases in size during the infection process [34,35,39]. We then cloned the *lacI* gene under control of the constitutive P_pen_ promoter into our four PALM-compatible vectors (pIA027, pIA028, pIA033 and pIA034), fused to the N-terminus end of the FP. To this end, the plasmids were linearized by digestion with the BamHI and NotI high fidelity enzymes (Fragment-I or ‘vector’) and assembled with a fragment containing the *lacI* gene, the RBS and the P_pen_ promoter (Fragment-II or ‘gene of interest’). The resulting four new plasmids pIA025 (*thrC::thrC-lacI-bsPAmCherry1*), pIA030 (*thrC::thrC-lacI-PAmCherry1*), pIA035 (*amyE::amyE-lacI-dronPA*) and pIA036 (*amyE::amyE-lacI-bsdronPA*) (Figure 2B and Table 2) were transformed into *B. subtilis*, generating strains RCL0937, RCL0938, RCL0939 and RCL0941, respectively (Table 1). These strains were then infected with phage SPP1*delX110lacO64* and imaged by PALM using HILO illumination to improve the signal to noise ratio of the bacterial cell thin samples. The previously reported strain GSY10004 expressing the LacI-mCherry fusion was used as a control of the localization of SPP1 DNA in infected *B. subtilis* cells. Monoinfected cells displayed a single phage DNA focus (Figure 3A) as previously reported [34,35]. Mild fixation of cells did not affect the localization pattern of the viral replication factories (Figure 3B). Infected cells expressing the LacI-PAmCherry1, LacI-bsPAmCherry1, LacI- dronPA or LacI-bsdronPA fusions were imaged both live (Figure 3C) and after fixation (Figure 3D). In all cases, successful localization of the bacteriophage DNA was observed in mono-infected cells. We are currently analyzing the lateral localization accuracies achieved by each FP fusion as well as the average number of molecules detected in the labelled LacI localizations. This quantitative analysis, which is unique to PALM, will allow to identify the number of molecules detected per cells/DNA cluster. It will also allow to quantitatively assess the codon-optimised versions of the FP relative to their non codon-optimised counterparts.

**Figure 3.**
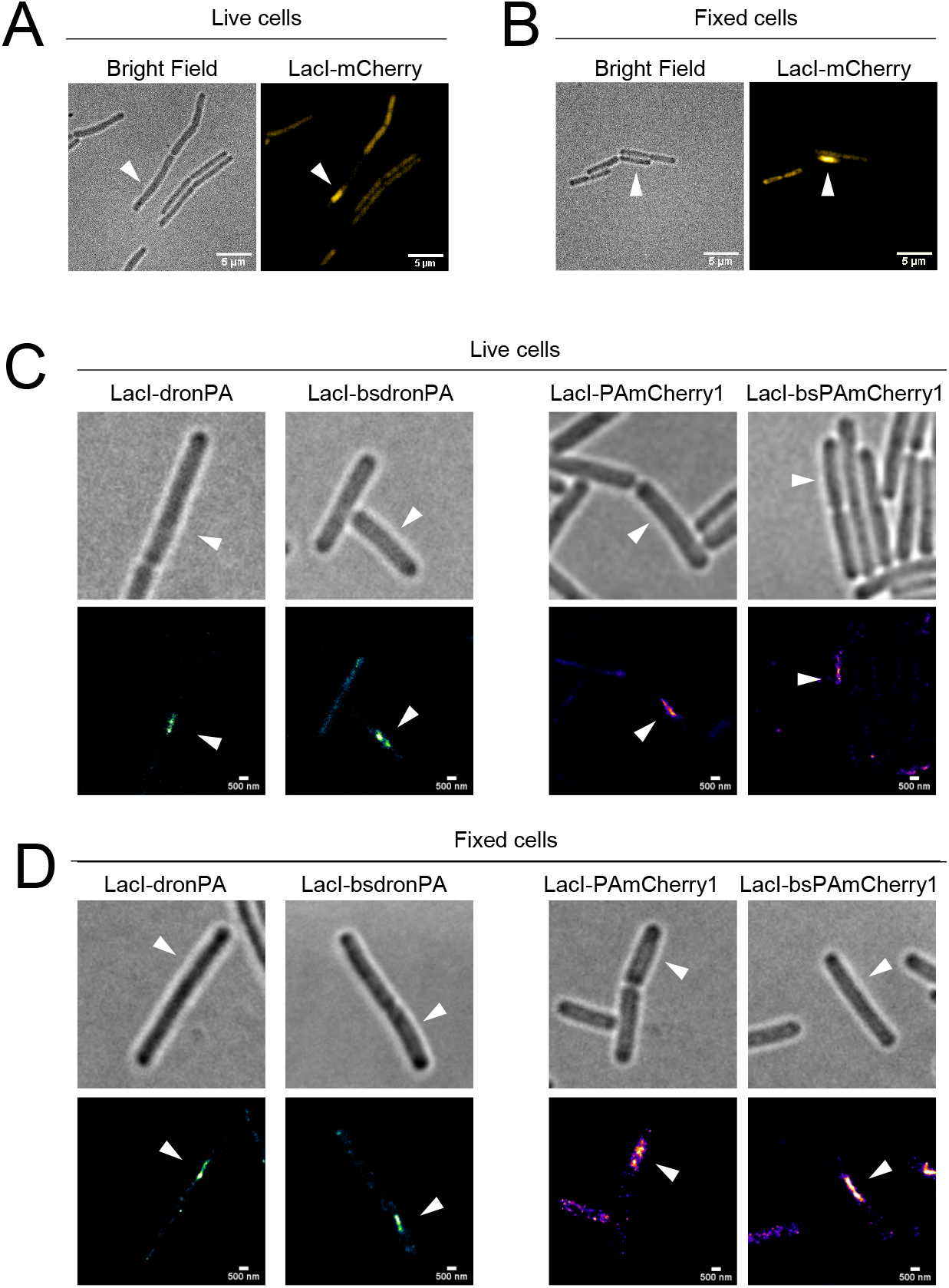
Visualization of bacteriophage SPP1 DNA in infected *B. subtilis* cells. Imaging of bacteriophage SPP1 viral DNA in infected *B. subtilis* cells for validation of our PALM-compatible vectors and confirmation of the methodology. *B. subtilis* cells constitutively expressing the different LacI-FP fusions from the ectopic loci were infected with phage SPP1*delX110lacO64* and imaged 50 min post-infection. In infected cells, the LacI repressor binds to the *lacO* operator repeats in the bacteriophage viral DNA, allowing visualization of the viral DNA. White arrows indicate the infected cells. **A-B**. LacI-mCherry localization in live (A) and fixed (B) cells of strain GSY10004. Scale bars, 5 μm. **C-D**. Single molecule localization of dronPA (strain RCL0937), bsdronPA (strain RCL0941), PAmCherry1 (strain RCL0938) and bsPAmCherry1 (strain RCL0939) fusions to LacI in live (C) and fixed (D) cells at 50 min p.i. dronPA/bsdronPA localizations are shown in green and PAmCherry1/bsPAmCherry1 localizations are in magenta. Scale bars, 500 nm.

## DISCUSSION

In this report, we describe the construction and testing of a set of eight integration vectors carrying the PALM-compatible dronPA and PAmCherry1 fluorescent proteins for use in *B. subtilis*. These vectors allow integration at the *amyE* and *thrC* chromosomal loci and can be used for SMLM studies of fluorescent fusions expressed under control of the promoters of choice, to widen the expression range and induction profiles. They also give the choice of the protein end to which the fluorescent protein can be fused. If combined in the same cell, they can be used for co-localization studies. Four of the vectors generated are dedicated to FROS LacI-*lacO* localization studies. Altogether, our vectors expand the toolbox for SMLM studies in *B. subtilis* and could be also used as templates for genetic engineering. All described plasmids are available through Addgene (catalog numbers pending) and could be of benefit to many researchers of the bacterial community.

## Data availability

The data generated during this study is available from an upon request from the corresponding author. All described plasmids and their complete sequences will be available through Addgene (catalog numbers pending).

## Acknowledgements

We thank Arnaud Chastanet, Magali Ventroux and Peggy Mervelet for technical suggestions. We thank Paulo Tavares for kindly providing strain YB886, plasmid pPT300 and phage SPP1 infection protocols and expertise, and for useful comments on the manuscript. This project was supported by funding from the Agence Nationale de la Recherche (ANR, grant BacVirRemodel to R.C.-L.) and from the European Research Council (ERC) under the Horizon 2020 research and innovation program (grant agreement No 772178, ERC consolidator grant to R.C.-L.).

## Authors information

Université Paris-Saclay, INRAE, AgroParisTech, Micalis Institute, 78350, Jouy-en-Josas, France Ipek Altinoglu, Rut Carballido López.

Abbelight, Cachan, France

Ipek Altinoglu

## Author contributions

Conceptualization and validation, I.A. and R.C.-L.; methodology, investigation and formal analysis, I.A.; resources, R.C.-L.; writing – original draft preparation, I.A; writing-reviewing and editing, I.A. and R.C.-L.; supervision and funding acquisition, R.C.-L.

## Corresponding Author

Rut Carballido López

